# Evolution of research in biomedical sciences - a network-based characterization based on PubMed

**DOI:** 10.1101/2020.09.10.291203

**Authors:** Karla Friedrichs, Michael Spranger, Sucheendra K. Palaniappan

## Abstract

The rapid growth of scientific publications every year makes it infeasible to keep pace with and survey manually, even for a specific field. Keeping up with literature and gaining a birds-eye view in a timely manner is crucial to the pursuit of scientific discovery and innovation. To help gain a clearer understanding of the state and progress of science and the nature of discovery, one can encode key information from these publications and represent them as a network. Observations on the structural evolution of these graphs can offer valuable insights on the dynamics at play. This work describes the construction and analyses the temporal evolution of a knowledge network of keywords (specifically focusing on genes/proteins, diseases and chemicals) from publications in the biomedical sciences domain. We compare and contrast the representations and evolution of these keyword networks types and find significant differences in the network growth, largely corresponding to our intuition. Furthermore, we focus on the formation and evolution of new links, which we argue corresponds to new scientific discoveries. Our findings suggest that these links are progressively formed in short network distance, leading to clusters of extensively studied keywords. This strategy, however, seems to impede ground-breaking innovation, which could be beneficial for research progress.

## Introduction

The rate of scientific publications has been constantly growing since the past century [1, 2]. But it is unclear to what extent this implies accelerating progress, as contributions made by research papers may differ significantly [3]. Previous work has shown that scientists often choose to pursue less ‘risky’ research directions that are promising positive results, but are unlikely to lead to great innovation [3, 4]. While this strategy might be advantageous for individual careers [4], it also has an impact on the way our knowledge expands. Our aim in this paper is to take a data driven unbiased approach to analyzing and understanding scientific literature from this point of view.

The amount of data generated by ongoing research has long become infeasible for manual investigation. Fortunately, several sources bring together large numbers of publications (among others, PubMed, Scopus, Google Scolar, SCIE) - although each has its limitations in coverage. Combining methods from big data, descriptive statistics and network science, the science of science (SciSci) aspires to make use of the available data for a meta-analysis on research [5].

To explore the dynamics of different fields, the biggest challenge often becomes representing the state of science in a meaningful way, ideally reflecting real-world events and research processes. A common approach to this task has been the analysis of networks formed by the total of these publications.

One of the first efforts has been made by Derek J. de Solla Price in 1965 [6], who drew conclusions about the practice of citing and the way papers are connected by analyzing inter-publication references on a large scale. Citation graphs have subsequently been used for a variety of tasks, e.g. examination of co-operations between cities and countries [7], consensus formation [8], or identification of scientific memes [9].

For the task of creating a taxonomy of science, Klavans and Boyack [10] recently reviewed three major types of citation networks: Direct citations (one document citing another links the two of them), as already considered by Price, have been used for a large-scale analysis only in 2012 [11], presumably due to technical limitations [10]. This method is judged the most accurate out of the three [10]. Co-citations (publications being cited by the same third paper are linked) have been a computationally more feasible approach throughout the early attempts to represent the network of scientific topics [12–14], but also appear in more recent studies working on larger data sets [15]. Finally, bibliographic coupling (documents citing the same third paper are linked), originally suggested by Kessler 1963 [16], has had a resurgence in the 2000s [17–19].

Combining the methods of direct citation and co-citation, emerging scientific topics can be detected in real time with few false positives [20], the review argues. Journal-based taxonomies, which are compared to document-based approaches in the same work, are judged less suitable to structure scientific knowledge [10].

Another approach considers researchers as nodes in a graph, creating co-authorship networks, for instance in order to compare collaboration conventions in different science fields [21] and on interpersonal, national, and international scale [22] or to analyse available expertise on neglected diseases [23].

It is debatable, however, to what extend papers, journals or even authors can represent knowledge. This question is addressed by Yi and Choi [24], who contrast citation networks with keyword networks. In the latter - depending on the field - substances and species, methods and theories, diseases and symptoms become nodes of the network, while edges signify various associations between the keywords.

They conclude that keyword networks, although not as straightforward in construction, can be a valuable alternative to citation graphs, since scientific terms clearly communicate elements of knowledge [24].

Similar co-word methods have been used for representations of diverse scientific fields, e.g. chemistry [25], neural network research [26] or scientometrics [27]. Su and Lee [28] applied tools of social network analysis to a keyword graph.

The evolution of keyword graphs, for instance, has been used to uncover different strategies for choosing research topics [4]. Another research direction performs link prediction on these networks, hypothesizing which keywords might be associated in future work [29, 30].

A noteworthy approach came from Shi, Foster and Evans [31], who constructed a multi-modal graph comprising both scientists and keywords (further split into chemicals, diseases, and methods) as nodes. Hyper-edges then represent single papers by connecting all researchers and topics involved. To gain an understanding on new links, it is argued, having researcher nodes is advantageous: In reality, how scientists choose research topics - which ultimately may lead to linkage - is significantly influenced by author-author relations. Researchers inspire each other, accept cooperation opportunities and, in general, are more aware of subjects discussed in their social proximity.

In this paper, we delve into the realm of biomedical science, more specifically, we aim to understand the developments in biomedical research focusing on analysing chemical, disease, and gene/protein keywords. For the specific goal of observing growth and cognitive potential of science, it has been argued before that the emergence and revision of scientific terms is a better indicator than mere publication numbers [32]. Therefore, we choose to construct a keyword graph out of crucial concepts in the given literature. Our goal is not only to find general research trends and differences between biomedical fields and reproduce them, but also to get insights on structural conditions that are likely to bear scientific innovation. At the same time, we undertake efforts to validate the keyword graph by matching research background or history of a selected keyword to observations made on the graph.

The aim of this work can be summarized as follows: In a descriptive approach, we find evidence that our method is a valid representation of scientific progress and get an impression of the dynamics at play. We then turn to the growth of the networks and ask in what places new links are formed and how this might impact scientific progress. The presented approach succeeds at capturing core intuitions about biomedical fields and motivates conclusions about research strategies that are in line with previous work.

## Materials and methods

### Dataset

For this work we selected PubMed [33] as our primary source of publication data, which contains over 30 million scientific abstracts [34]. We chose to focus on extracting and tagging 3 themes of concepts from these publications, namely chemicals, diseases, and finally, genes/proteins (summarized as ‘gene’ in the following). In terms of the timeline of these publications, we focused on publications ranging a time span of 50 years (leaving out abstracts published after 2017 to account for possible delays in updating PubMed, which could lead to wrong inference from our analysis). The full data set was downloaded and obtained from the bulk downloading service of the National Library of Medicine data distribution service. It is to be noted that MEDLINE started its record in 1966, but later included almost 60 thousand noteworthy papers previously published [35]. The pre-1966 data therefore has to be seen with a grain of salt, as is might be biased by the editors’ choice.

Our process of collecting and preparing data as well as the construction of our keyword graphs is summarized in Fig 1 and largely follows the approach taken by [3]. Specifically we first analyzed the complete PubMed literature for overall trends, and then focus on a specific subgroup to drive our findings after observing that the larger trends stay intact in this subset.

**Fig 1.**
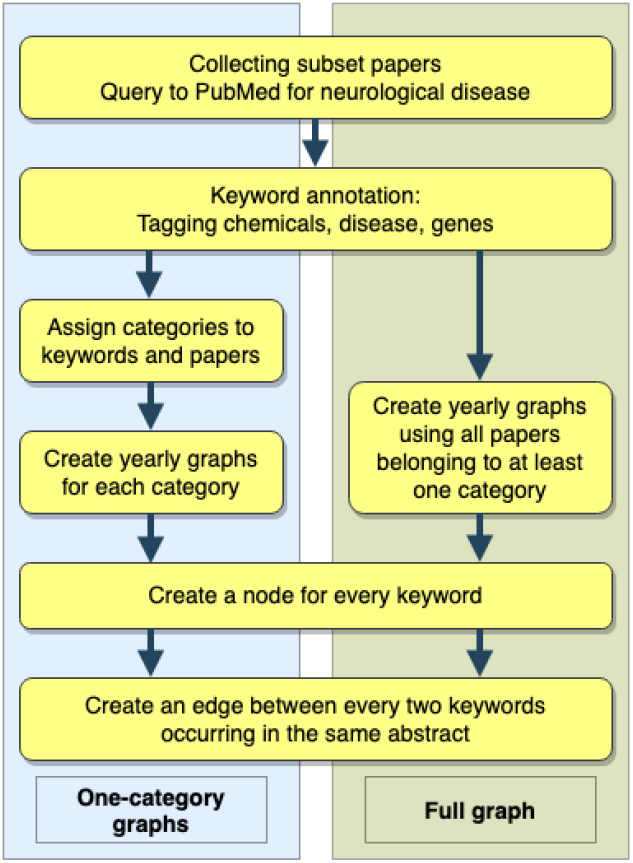
Process of collecting data, labeling keywords, and constructing the keyword graphs.

Our publication subset focuses on the nervous systems, which has seen a lot of interest in the research community recently and would be a good showcase for our inferences. Only publications containing at least one chemical, disease, or gene keyword were included, leaving us with 2.1 million abstracts. This not only reduced the computation efforts, since the main part of our work concentrates on a keyword network and requires expensive calculations, but also made it more feasible to keep an overview of relevant influences on the incoming data.

The following search term was used to extract our subset from the PubMed database:

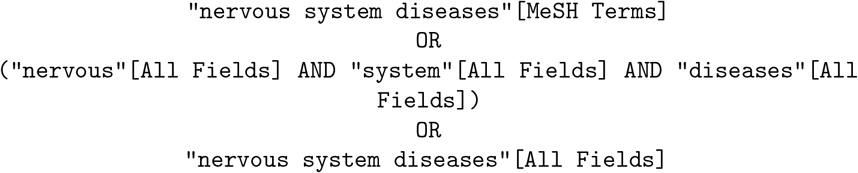

The papers come with keyword annotations - central concepts in each article have been manually indexed [36] with uniform keyword identifiers. In many cases, however, rather general tags were chosen for specific terms. As the annotation quality is a crucial factor for the outcome of our analysis, we decided not to rely on the existing annotation. Instead, we run every article through a named entity recognizer for tagging keywords. For annotating concepts we rely on the Pubtator [37] service which uses their pre-trained suite of named entity recognition tools including TaggerOne, GenNorm for gene normalization, DNorm for diseases, and tmChem for chemicals.

Given an abstract, Pubtator returns the keywords/concepts in the abstract with their identifier. In our work, we consider three keyword vocabularies: diseases are assigned Medical Subject Headings (MeSH) [38], chemicals and drugs are matched to either MeSH identifiers or Chemical Entities of Biological Interest (ChEBI) [39], and finally NCBI gene identifiers [40] are used to tag genes.

The downloaded files were parsed and imported into a database where the meta information and abstract content is stored. Details on the composition of the database can be found in S1 Table.

## Construction of a co-occurrence network

In our keyword network each node is a research subject and edges represent relations known to hold between these concepts. More precisely, network nodes are created for each keyword identifier appearing in our corpus, and connections are extracted from the papers’ titles and abstracts.

Here, as a notion of relatedness, we simply assume a link to exist if two keywords have been mentioned in the same abstract, since an abstract is expected to contain only a concise range of concepts relevant to the paper’s thesis. This presumption is discussed in more detail at the end of this paper.

Fig 2 illustrates how a part of the graph could look like. The depicted terms each get their own node, as they can be assigned distinct MeSH identifiers. As an example, a paper describing the case of a patient with Broca’s aphasia might use the keywords *Broca’s aphasia, stroke*, and *brain ischemia*, connecting each of the nodes. A second paper might have associated *diabetes* and *stroke*, creating paths from *diabetes* to the other three keywords.

**Fig 2.**
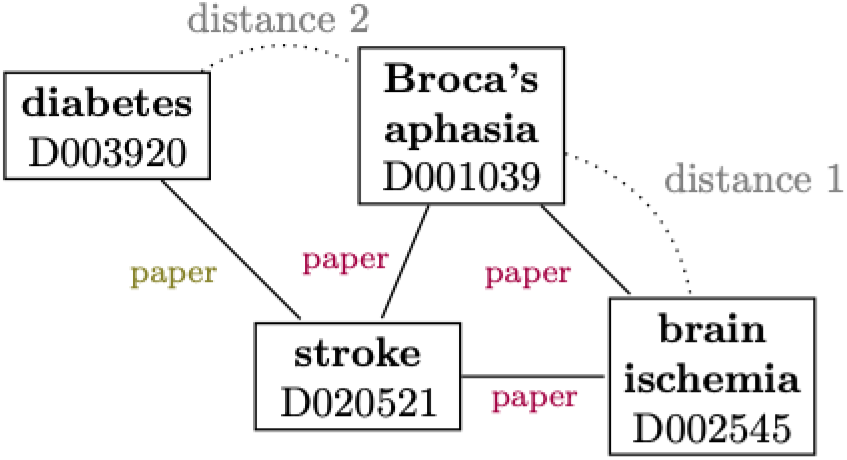
Constructed example of a substructure that could be found in the keyword network.

## The co-occurrence network

We also point out that directly linked nodes are in distance 1, since the shortest connecting path consists of exactly one edge, while nodes sharing a neighbor have a distance of 2. This concept will be of high importance later in our analysis.

First, a graph is built including all publications up to (inclusively) 1947. Subsequently, we created snapshots for every year up until 2017, incorporating new findings as they come up.

Each graph is meant to capture the state of biomedical research at a certain point of time: nodes are added as new genes, diseases or chemicals are discovered, edges are established when two concepts are associated for the first time.

## Results and discussion

### Comparing scientific progress in three biomedical fields

Splitting the keyword vocabulary into chemicals, diseases and genes allows for their individual characterization and shows a surprisingly different progression of the corresponding networks. Primarily, our aim is to outline the sharp contrast between the disease and gene keywords, as the chemical-only graph seems to lie in between the other categories in most of the examined properties (see S2 Appendix for some observations). We observe clearly different trends (in Fig 3) in the keyword usage and evolution, and subsequently check if our graph captures publications on diseases and genes in varying states of research.

**Fig 3.**
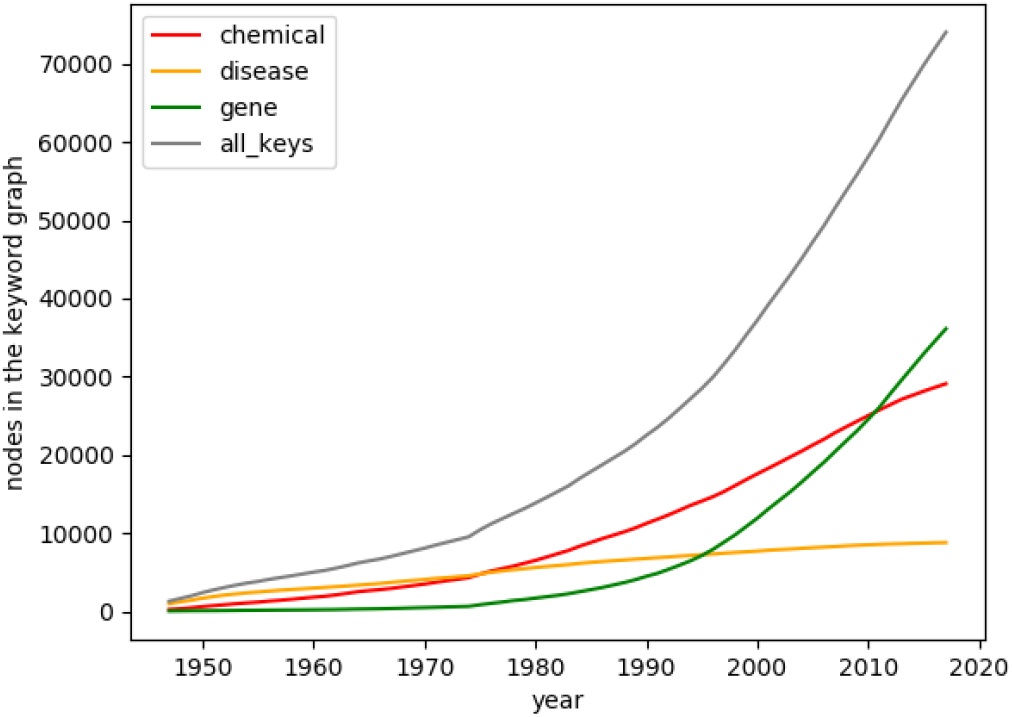
Graph sizes. The number of nodes in the keyword graphs is depicted for every year.

Disease research seems to focus on a small and slowly expanding set of keywords - in 1947, it starts as the biggest network, but accumulates a total of little more than 10,000 keywords until 2017. On the other hand, close to no genes appear up until the mid-70s, after which the graph grows up to a size of more than 35,000 nodes. The first nucleic acid sequencing in 1965 and the evolution of genomics techniques during the 1970s [41] heralded the accelerating publication rate during the 1980s and 1990s, when amongst others the human genome project was launched [41], events that are made visible by the network growth.

One might hypothesize here that the graph sizes simply correlate with the number of publications about diseases, chemicals, or genes. However, Fig 4 shows how both publication rates and the count of keyword occurrences are considerably higher in the disease category, underlining the discrepancy between disease and gene papers in respect to generating new keywords.

**Fig 4.**
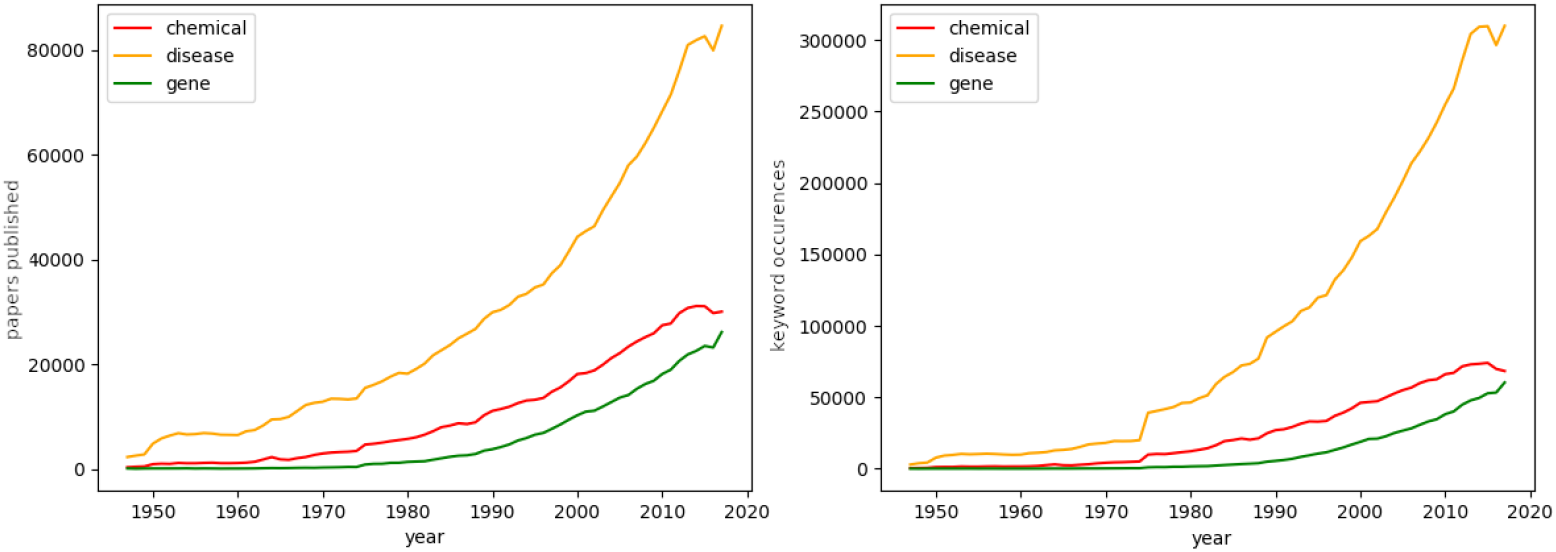
Publications and keywords per year. Left: Publications add to the number of a category if their abstract contains at least one corresponding keyword. Details on the dataset are found in S1 Table. Right: Total keyword occurrences. Keywords are counted more than once a year if they are mentioned by multiple papers. Up until 1975, there are at most 250 keywords used in genomics per year.

As the corpus was retrieved by a query for papers about neurological diseases, it is no surprise that a majority of the publications contains disease keywords. In fact, from the set of 2.1 million papers, only about 60,000 do not belong to the disease category (S1 Table). For gene publications, the figures again show growing rates roughly corresponding to the historical emergence of genetics, but the number of articles in this field stays the smallest of our subset by far.

The publication rates alone apparently cannot explain why gene keywords surpass the other categories in number and a major part of the articles forms the smallest network. We propose two main reasons here, one inherent to the scientific fields, the other based on the progress of research.

First, disease keywords stand for symptoms and syndromes, which motivates an intuitive explanation: diseases are complex and consist of multiple causative elements, including genes/protein malfunctions, and generally the focus is on developing treatments or investigating causes to known illnesses (see S1 Appendix for more details) which are long horizon. In contrast, with the explosion of genomics and high throughput technologies the focus has been on discovering more genes.

The next section will elaborate on how the ongoing research on well-known diseases shapes the keyword network. Here, we want to illustrate once more the different roles new keywords seem to take in the gene and disease research. Fig 5 shows what amount of novel concepts there is in all keyword occurrences - to put it differently, if a keyword is found in a paper abstract, what are the chances it has never appeared before?

**Fig 5.**
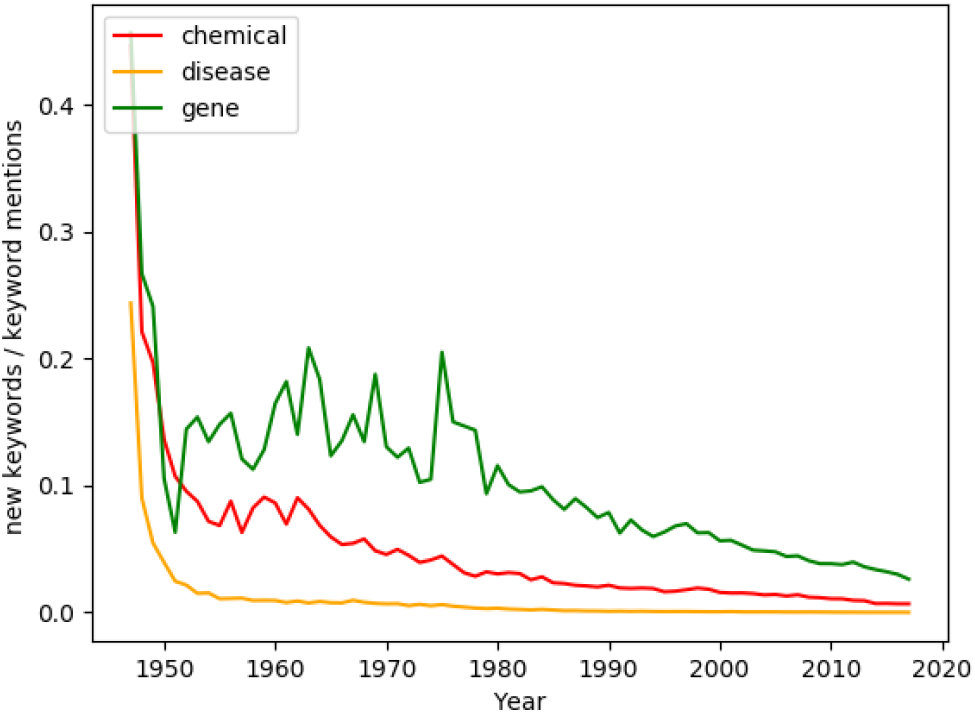
Significance of keyword introductions. The ratio of keywords newly introduced to the total number of keywords mentioned in a year (Fig 4.). The latter is not to be confused to the number of unique keywords. A keyword is counted as ‘new’ only once in the year it first appears.

As expected, the rate of new disease keywords stays close to zero from the 50s on: Most of the time, scientist appear to publish about known diseases instead of examining new symptoms.

For gene research, the network before the 70s is extremely small, making the rate of new keywords less meaningful. In the 80s, every tenth appearance of a keyword is, in fact, describing a gene not sequenced before. This rate remains higher than in the other categories, but drops to below 5% after 2005. Keeping in mind that in the year of introduction, a keyword is counted as new only once if mentioned by multiple abstracts, this still suggests a high importance of gene sequencing. We would like to point to S1 Fig for an impression of the rate at which keywords are discovered.

This observation also leads to our second hypothesis about the differences between disease and gene keywords. As mentioned above, genomics has only been a notable research direction for about 50 years, which makes it a relatively young field. We credit the high number of new keywords partially to the early state genomics is still in. Fig 5 - as well as graphs on the network evolution further below - suggests the gene network might eventually fall in line with the ‘older’ sciences. It seems plausible that sequencing projects will eventually become less important, as our knowledge of existing genomes becomes more complete.

## Sparking scientific innovation

A keyword graph is our attempt to grasp how crucial concepts are connected through research. The preceding analysis on the keyword distribution offers an informative first view on biomedical research. But science is more than just a sequence of publications - findings build on previous findings, papers inspire new research and there is active exchange in the scientific community.

Scientific innovation, in this paradigm, is represented by a new link between two keywords, based on a paper associating the concepts for the first time. In the following, we have a closer look at the structural evolution of the keyword graph, as this might hint at our main question: How does science innovate?

In general, keywords have direct links only to a very small set of other keywords, compared to the overall size of the graph. Consequently, the keyword graphs are highly incomplete, as demonstrated by a low density. We give a definition of density for a graph G = (V,E) in Eq (1):

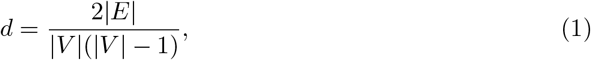

where E is the set of edges and V the set of vertexes.

The highest score was found for the disease graph and amounts to little more than 0.03, while chemicals, genes as well as the complete graph have densities between 0.001 and 0.003 (Fig 6). Intuitively, the scores can be translated into the percentage of other keywords an average keyword is connected to. In 2017, the disease network has 9000 keywords and an average degree of almost 300 - the gene network on the other hand has 36,000 keywords with a mean of only 25 connections, resulting in a very low density.

**Fig 6.**
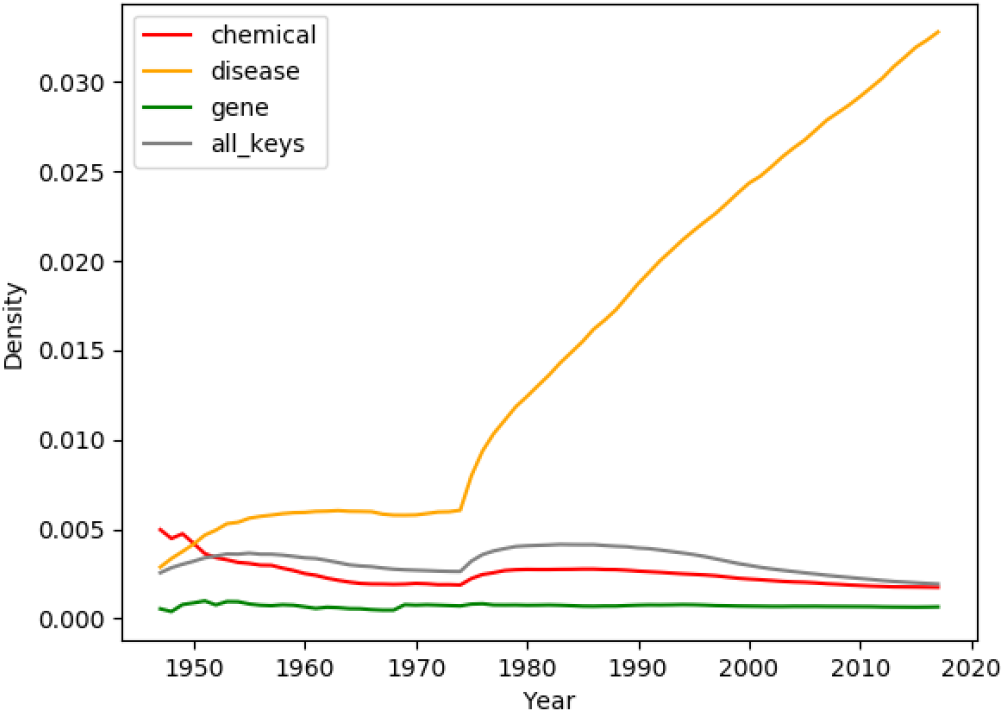
Density. Density of the three category graphs and the full graph as defined in Eq (1) A complete graph has a density of 1, the density is 0 for a graph consisting of isolated nodes (keywords) only.

Two keywords having a known relation is rare - which underlines the question of how scientists choose their research subjects even more.

The existing links, in fact, are not evenly distributed throughout the network. Instead, keywords are typically embedded into a strongly interacting neighborhood. Evidence is provided by the average clustering coefficients. If a term has a coefficient of 1, this means that all keywords co-appearing with this terms have pairwise co-appearances themselves.

In 2017, the clustering coefficient is at 0.75 for the full network and still around 0.7 and 0.65 for diseases and chemicals, respectively (Fig 7). The gene-only graph shows the least clustering with a coefficient of slightly over 0.5 (Fig 7), which conforms to the high number of new keywords introduced each year.

**Fig 7.**
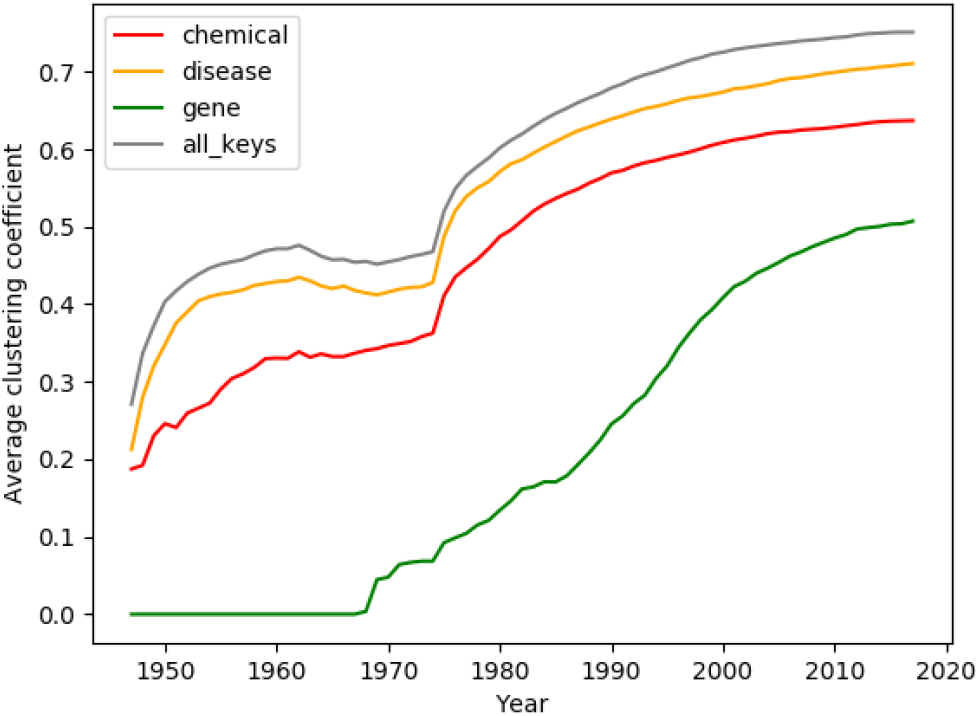
Average clustering coefficient. The clustering coefficient of a node is defined as the ratio of existing node triplets to the number of possible node triplets in its direct neighborhood, where a node triplet means two neighbors of the center node being connected to each other. The figure displays the clustering coefficient averaged over all nodes of a graph.

This observation is a first hint to our initial problem. Of course, this connectivity is fueled by our method: An abstract mentioning a set of keywords immediately fully connects them to each other. However, the average degree is much higher than the mean number of keywords per paper in all of the graphs (S3 Fig) - this means the clustering cannot be explained by large numbers of keywords being interconnected by single papers alone.

Instead, we suspect it also originates from a great number of connections between already close keywords. This is illustrated in Fig 8, which describes the three-category keyword graph: Apart from the first years, when the graphs are still only weakly connected, and especially from 1980 until today, new links are mostly established between keywords that were already close. Specifically, the vast majority of connections are made between concepts that are connected by a path of length 2, meaning there is a common third keyword they both co-appear with in another abstract. The same tendencies can be observed for all graphs, although the distance-2-discoveries start to outweigh other connections later in the chemical and gene networks (S4 Fig, S5 Fig, Fig 10).

**Fig 8.**
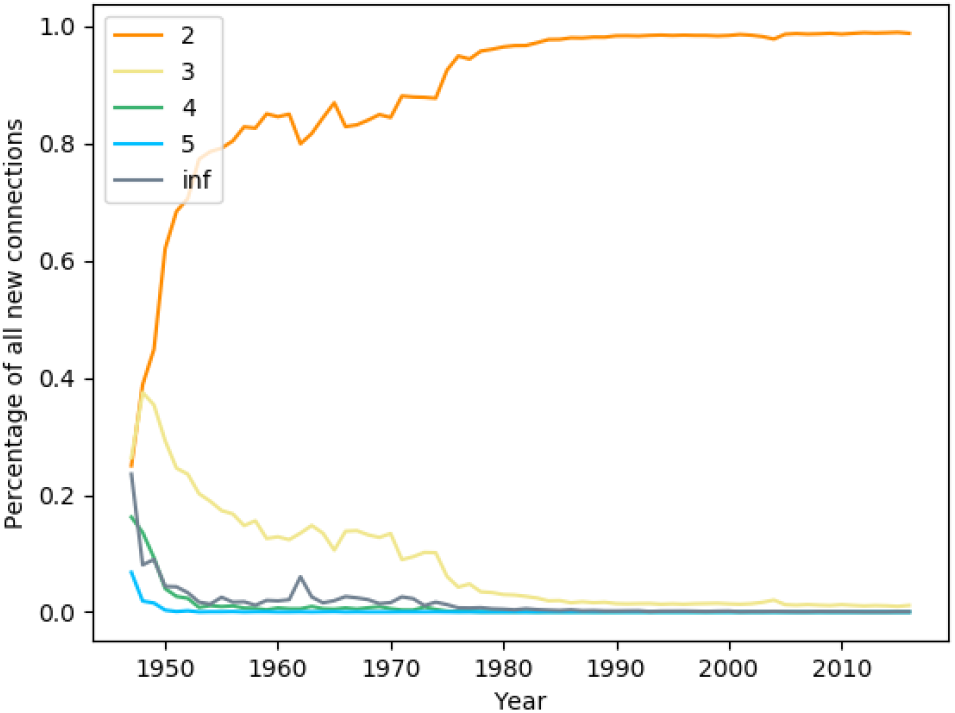
Previous shortest path between connected keywords. For each new keyword connection, the preceding year’s graph was searched for the shortest path between the nodes using Dijkstra’s algorithm. ‘inf’ signifies that the nodes had no connecting path.

**Fig 9.**
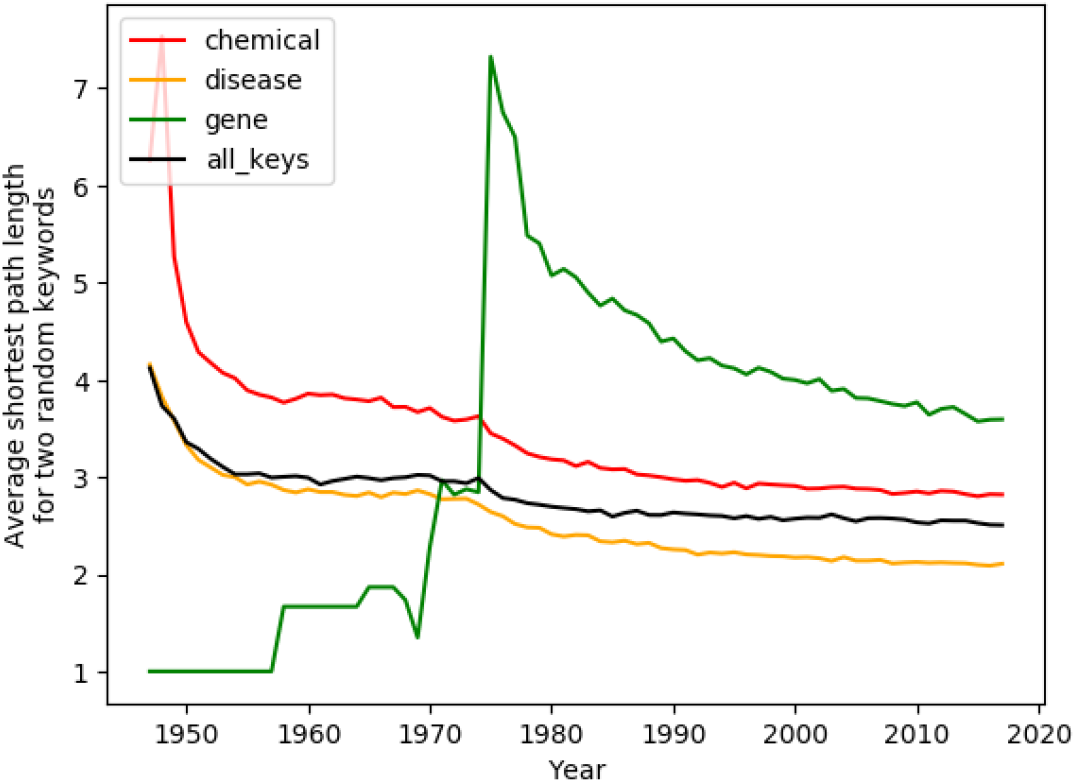
Average length of the shortest path between two keywords. Computing the average shortest path between two keywords turned out to be infeasible, as the full graph contains at times more than 300.000 keywords. Instead 100.000 node pairs were selected randomly for years surpassing this number of possible keyword combinations.

**Fig 10.**
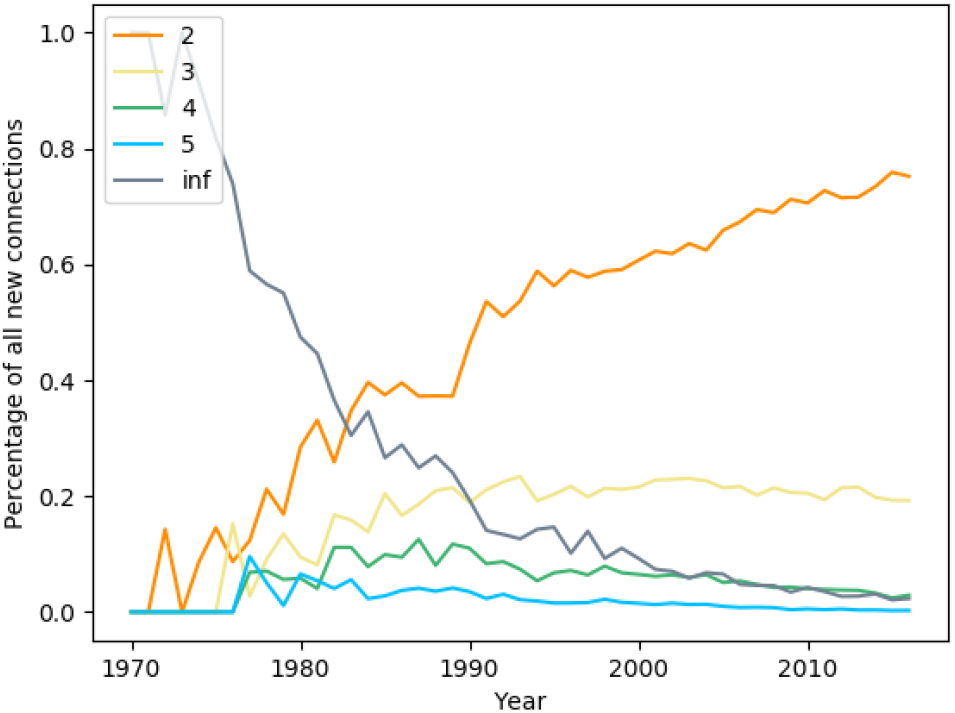
Gene network: Previous shortest path between connected keywords. For each new keyword connection, the preceding year’s graph was searched for the shortest path between the nodes using Dijkstra’s algorithm. ‘inf’ signifies that the nodes had no connecting path.

Next we attempted to analyse whether this overwhelming predominance of short distance attachment is due to short paths between keywords in general - or whether it can be interpreted as part of the scientific process. Indeed, since the 1980s already, the average path between two nodes in the complete graph is little longer than 2.5 9, and manual investigation showed that mainly paths of length 2 or 3 and a few outliers of length 4 were sampled.

While Fig 8 does indicate a preference towards connecting close concepts (e.g. those with a common neighbor), it is to be noted the recent nature of the keyword graph does not allow for far-stretched discoveries anymore. We therefore turn our attention to the gene graph: again, it shows similar tendencies as the others but seems to be in an earlier state of development, which allows for more interesting insights into the formation of new links.

Just like the publication and keyword rates before, the network properties appear to make the birth of genomics visible. During the initial years, the graph offers little valuable information, since the network then consists of only a few genes. Consequently, new links are almost exclusively connecting isolated graph constituents.

In 1980, half of the new connections are still established between nodes of different graph components, 30% between nodes in distance 2, and 10% in distance 3 (Fig 10). Finally, during the 1990s, the graph resembles more to the other fields with the rate of distance 2 connections surpassing 60 % and the percentage of links between unconnected nodes falling below 10% (Fig 10). Even today, genes are on average still in a relatively high distance - about 3.6 edges in 2017 (Fig 9).

In the other graphs, average path lengths between 2.5 and 3 make close attachment less of a surprising observation. The gene network, however, provides stronger evidence: Even though compared to chemicals and diseases there are less connections in distance 2, these short distance links still clearly predominate with 75% in 2017 along with 20% links being made in distance 3 (Fig 10).

These findings conform to related projects. Shi, Foster and Evans observed a preference for short-distance attachment in their multi-model graph and conclude:

> Taken together, these findings suggest that when a scientist chooses a new topic to study or adopts a new method for her investigation, she is highly likely to choose something directly related to her current expertise (or something used by a collaborator). [31]

We would like to expand this explanation. Not only does it seem likely that scientists pick up subjects bordering their past research, this dynamic also fits a well-known model of innovation.

The term *adjacent possible* [42, 43] *-* put simply - describes the undiscovered knowledge that lies just next to what has been anticipated, and it suggests that each innovation clears the way for new insights. Loreto et al. [44] developed a mathematical model of this idea; they picture scientific knowledge as a subgraph of a huge network *(the actual*), around which lies the undiscovered (the *possible*). An innovation joins a bordering node to the knowledge, thereby expanding the edge and paving the way to new discoveries that had been unreachable before.

Arguably, linking two keywords in distance 2 in our graph corresponds to the *adjacent possible*. There is less of a mental leap if the keywords have been used in connection with the same third term, whereas more creativity might be necessary to associate keyword without a common context.

In a lot of cases, the distance between keywords gradually decreases before they ultimately get connected. In Fig 11, we made an attempt at visualizing this dynamic. Of the keywords that co-appear for the first time in an abstract published in 2016, 100,000 pairs were chosen at random. Then, in 10-year-steps from 1947 on, their relation was observed, discriminating between those keywords that have not yet appeared in our data (‘not in graph’), those that did appear, but are located in separate components of the graph, and those that already have a connection through other nodes.

**Fig 11.**
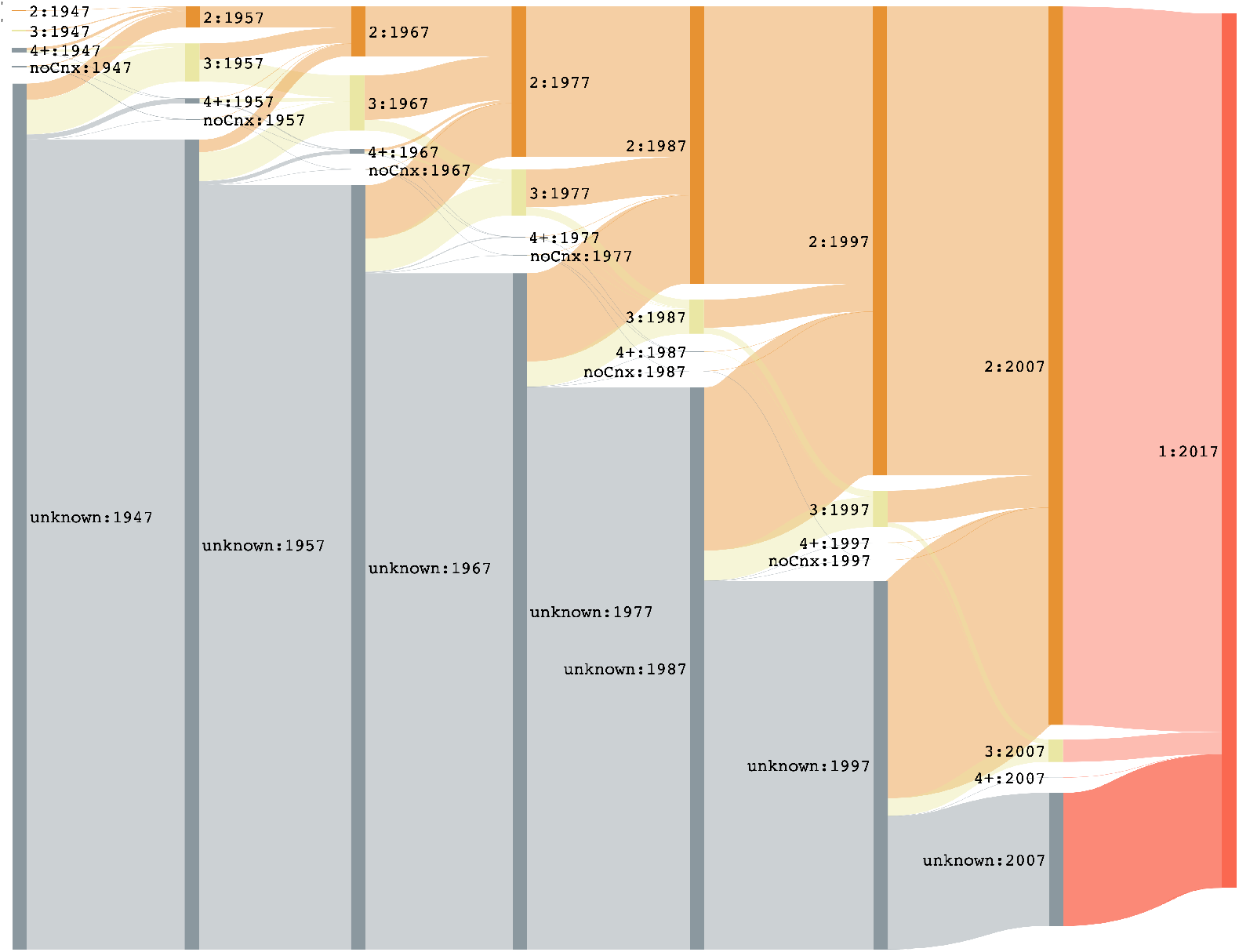
Observing keywords that co-occur in 2016 throughout time. Of the new links formed in 2016, 100,000 were chosen at random. In 10 year steps, the ‘relation’ of each pair is observed. A number denotes the length of the shortest path connecting the keywords, *‘not in graph’* means at least one keyword has not appeared in any abstract yet, and *‘unconnected’* signifies both keywords have appeared, but there is no connecting path.

Although the 10-year-steps, which we used for better readability, might mask some of the more fine-grained dynamics, there is an interesting observation: While many pairs go from ‘not in graph’ to having a path of length 2, a few first have a distance of 3 or more steps, which often decreases to 2 in subsequent timestamps.

According to the hypothesis of the *adjacent possible*, lowering the distance in the graph between the latter pairs might have been an important step for ultimately discovering their relatedness.

On the other hand, the rare links between keywords in long distance often represent innovative, rewarded, and possibly impactful discoveries, according to related work on similar networks of molecules [3, 4] as well as co-citation networks [45]. A manual investigation of long-distance links sampled from our network conducted by experts seems to support this hypothesis. Rzhetsky et al.([3]) even simulate different research strategies on keywords networks and find that increased work on these ‘risky’ long-distance connections might prove advantageous for scientific progress.

We would like to illustrate the idea of ‘innovative’ links by looking at an example. The keywords *sunburn* and *indomethacin* become eventually connected. Indomethacin is a non-steroidal anti-inflammatory drug which was traditionally used for curing Gout, Rheumatism, Arthritis etc. and it works by blocking the production of substances that cause inflammation. It was interesting to note that the use of indomethacin solution for sun burnt skin of humans resulted in marked decrease in UV light induced erythema. The effect of indomethacin for skin burn was not an obvious hypothesis until the 1975. Probing the shortest connection between these two keywords prior to 1975 resulted in the path illustrated in Fig 12.

**Fig 12.**
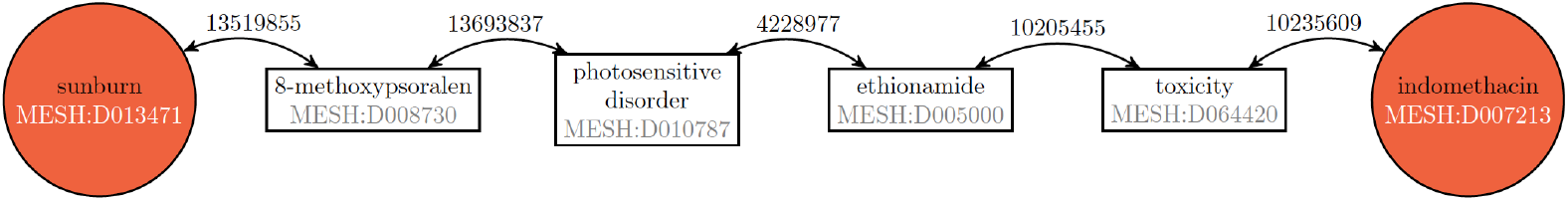
Shortest path *sunburn - indomethacin* in 1974. The keywords are linked in 1975. For each intermediate connection, the PMID of a paper supporting the relation is given.

It is interesting to note that 8-MOP was used traditionally in the context of sunburn, and generally for photosensitive disorders. Indomethacin on the other hand was considered for a general class of photosensitive disorders resulting in the indirect connection between indomethacin for use in sunburn.

Similarly one interesting example of keywords which were 3 steps apart but which got eventually connected in 1982 was that of “Alzheimer’s disease” and “interferon-gamma”. It was not known until then that there was an impaired interferon-gamma production in leucocytes for people with Alzheimer’s disease. In fact since then there is a lot of research which has proven the correlation between cytokine secretion and disease state in Alzheimer’s.

To summarize, the PubMed papers create a small-world graph of keywords: Most keyword pairs are connected in only a small number of steps, even though the average degree of each node is low compared to the full keyword set. In a small neighborhood, concepts seem to be strongly related. The majority of new edges is being established between keywords that share a common neighbor - and not only in further developed graphs with very short paths between all keywords - which effectively creates a node triangle and thus increases the clustering.

## Limitations

The presented results largely depend on the idea that keywords are related as soon as they appear in the abstract of the same paper. While this approach offers an easily implementable graph and link analysis, the results have to be seen with a grain of salt.

Considering that some abstracts are rather long or contain ten different keywords, co-occurrence can be a weak definition. Concepts might get a link in the graph although they appear far from each other in the abstract and are completely unrelated. Furthermore, the abstracts are in no way linguistically processed, so negation is not detected and papers stating that two concepts are *not* related will still produce an edge.

It has to manually evaluated how much noise our connection assumption actually introduces. In S4 Appendix, we have a look at an example abstract and discuss the edges it establishes.

To prevent negative statements and coincidental co-occurrences introducing faulty edges to the keyword graph, we suggest two alternative assumptions. Firstly, a straightforward refinement to the given definition would be to only include edges surpassing a threshold of papers that make the same connection. Meaningful co-appearances of concepts are expected to be repeated in multiple publications, unlike more random links. Upon manual comparison of these ‘reinforced’ edges and links made by only one paper, this approach seemed highly promising and should be taken into consideration in future work.

However, there is a risk of losing an advantage of the automated knowledge graph generation. By concentrating on concept relations frequently mentioned in literature, discoveries that attracted less attention will stay as unnoticed as they would be by a human.

Secondly, the links between keywords appearing in the same abstract could be refined by a notion of linguistic dependency. While syntactic parsing potentially offers a more precise judgement on constituency relations, even a simple measure like linear distance in text might improve the overall link quality.

Nonetheless, even if two keywords appearing in the same abstract is not an infallible argument for a real-world connection, we would like to note that the opposite still holds: a pair *not* co-occurring even once is a strong indicator that no relation has been found so far - given that the publication database is fairly exhaustive.

## Conclusion

In this paper, we made an attempt to analyse scientific progress as represented by a growing network of key concepts. We built on a rather loose definition of relatedness of concepts, which introduced noise to the network, masking smaller details. However, it allowed for the automatic construction of huge networks with ten thousands of keywords from biomedicine. These graphs did not only show characteristic traits of different biomedical subfields, but also were in line with related work representing science by aspects other than keywords.

Finally, we characterized the network evolution and found that in the discussed areas of research, there is a tendency of forming ‘clusters’: groups of concepts are frequently associated among each other, but there is little work examining more far-stretched relations. While this observation can be regarded as the natural way of knowledge expansion, as summarized in the theory of the *adjacent possible*, we also point to related work that has shown risk-taking in the choice of research subjects might prove beneficial for scientific progress.

## Supporting information

**S1 Fig.**
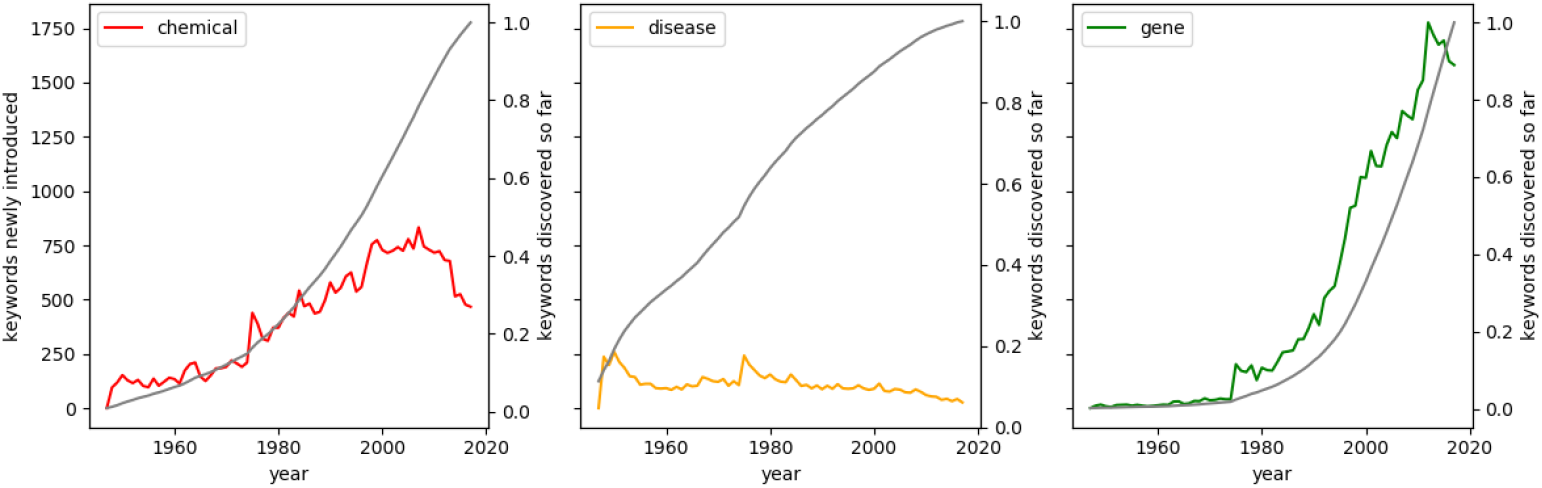
New keywords and vocabulary progress. The number of distinct keywords mentioned for the first time in the corpus is depicted on the left axis. The right axis shows the percentage of the 2017’s keyword set that has already been introduced in a certain year.

**S2 Fig.**
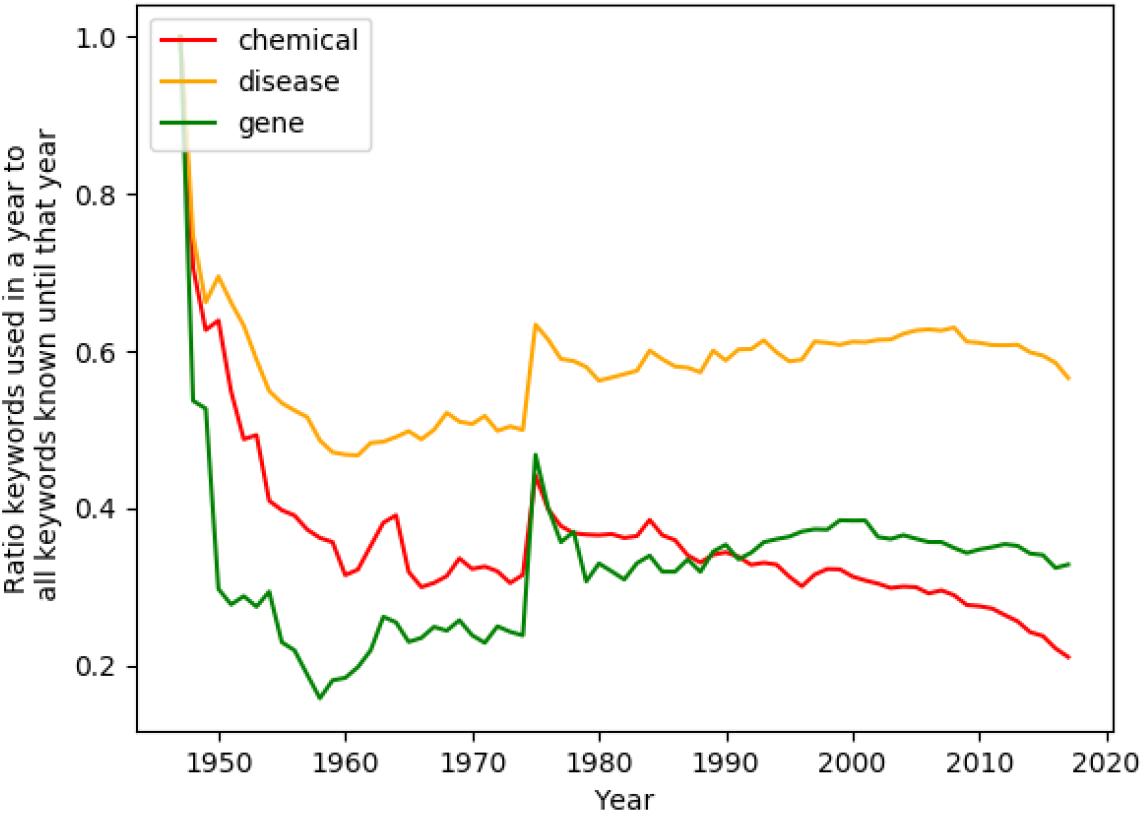
Percentage of known keywords reused in a time span. Each data point shows the ratio of keywords (known up to the corresponding year) that were mentioned at least once by a paper during that year.

**S3 Fig.**
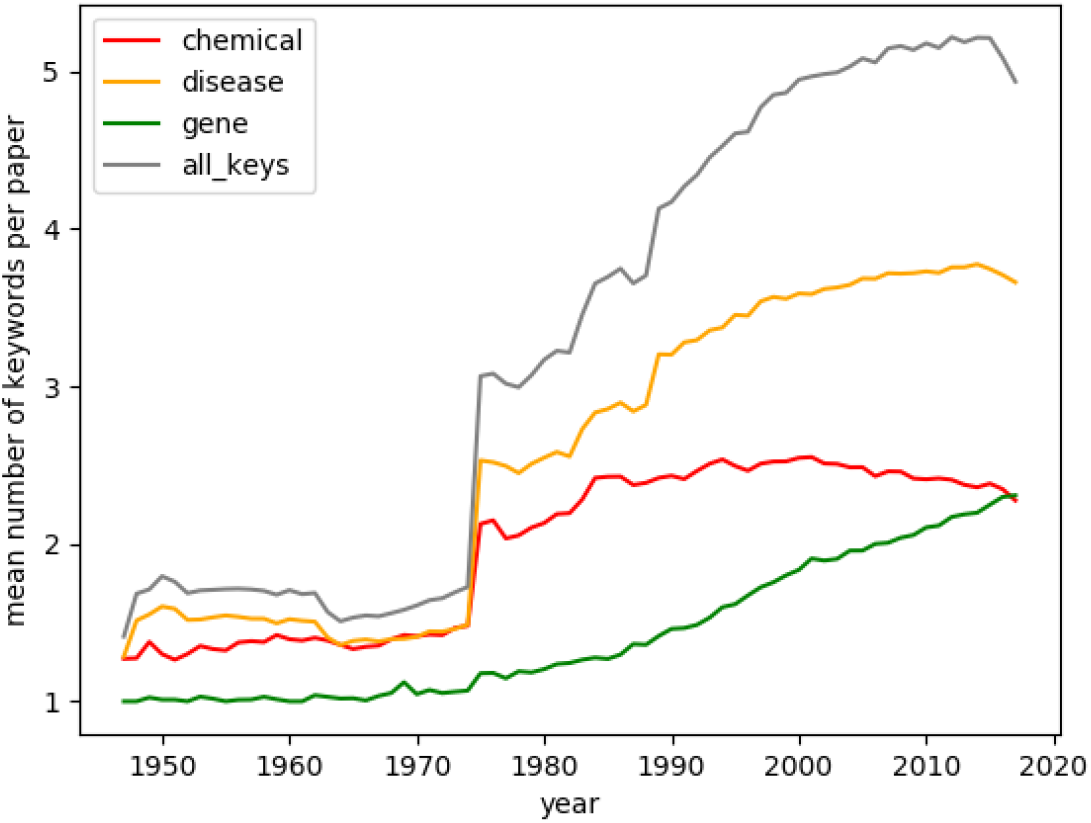
Average number of keywords in an abstract. For each paper, the unique keywords (keywords with distinct identifiers) were counted. The numbers were then summed up in each category and divided by the number of papers belonging to the respective category.

**S4 Fig.**
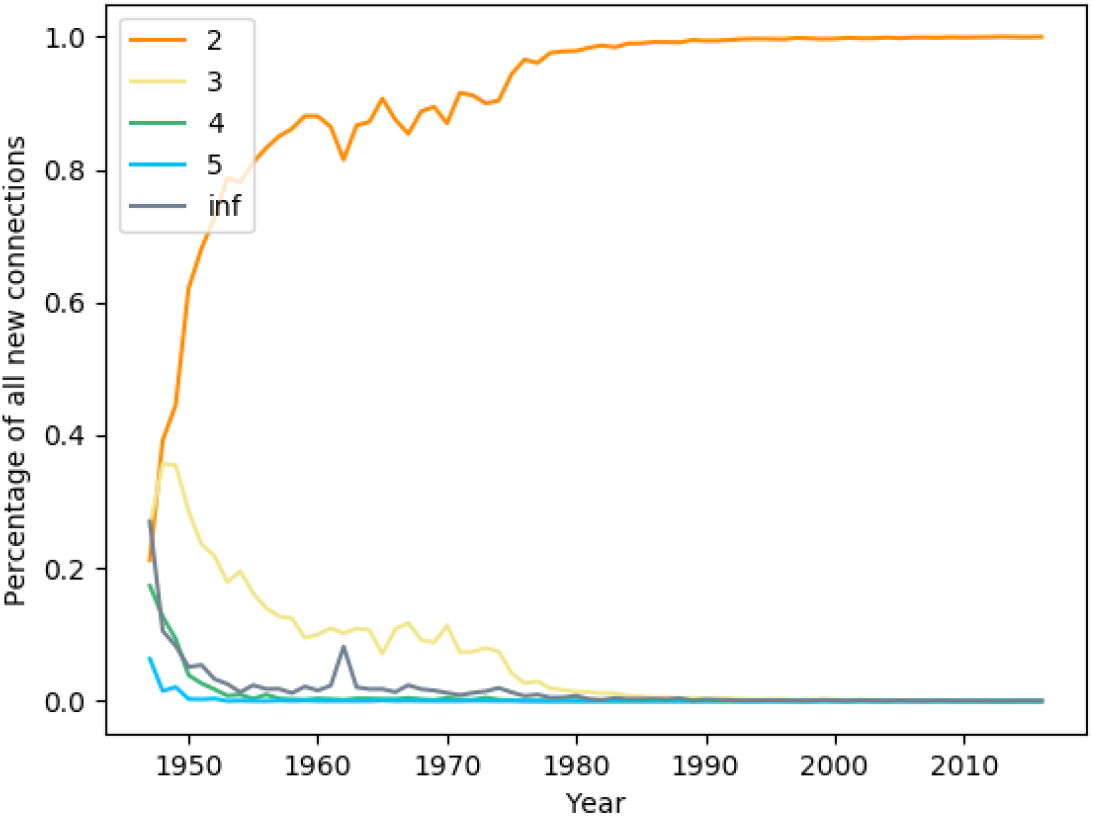
Disease network: Previous shortest path between connected keywords. For each new keyword connection, the preceding year’s graph was searched for the shortest path between the nodes using Dijkstra’s algorithm. ‘inf’ signifies that the nodes had no connecting path.

**S5 Fig.**
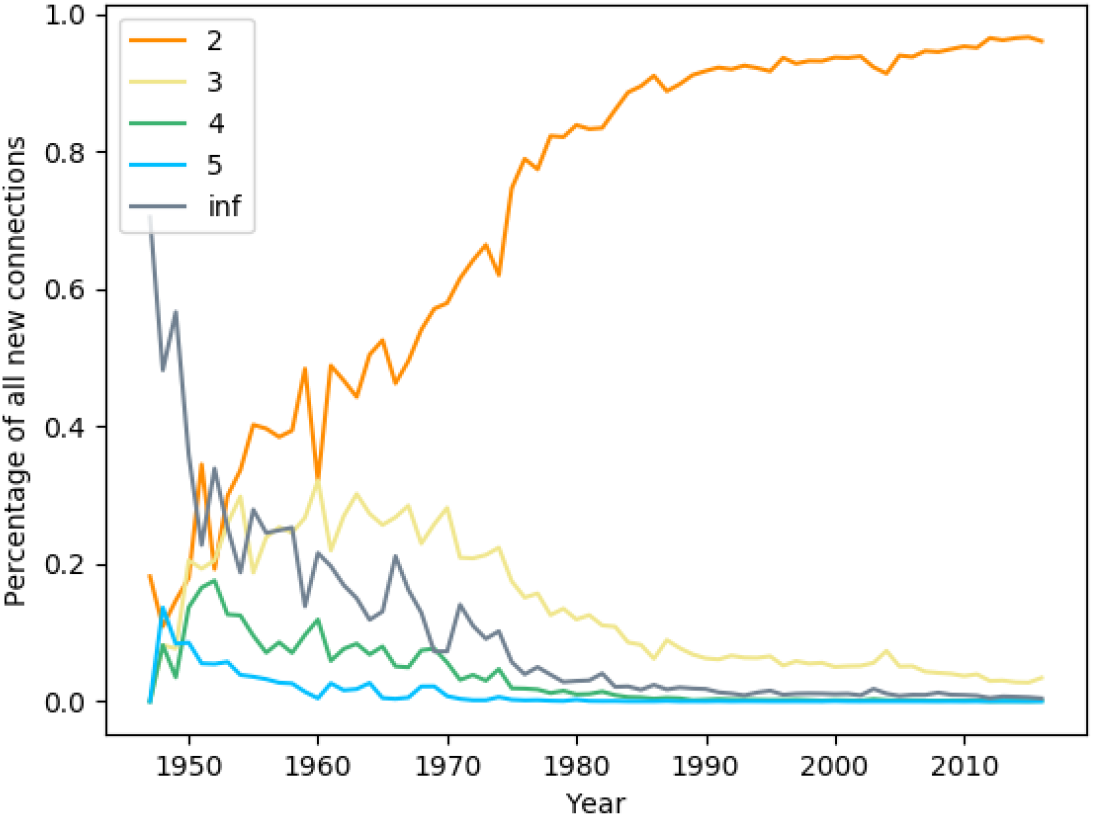
Chemical network: Previous shortest path between connected keywords. For each new keyword connection, the preceding year’s graph was searched for the shortest path between the nodes using Dijkstra’s algorithm. ‘inf’ signifies that the nodes had no connecting path.

### S1 Appendix. Further information on the disease network

Above, we have noted that new concepts are introduced to the disease network only sparingly. In fact, many keywords have been in use since the beginning of record in our database.

S1 Fig shows that around 1955, more than 30% of the keywords known until today have already been mentioned. The remaining keywords are gradually added with only around a hundred new keywords being introduced every year.

Not only does the field apparently focus on ‘known’ symptoms, but it also consistently covers large parts of the existing vocabulary in ongoing research. Looking just at the publications of a single year, around 60% of all terms known until then appear at least once (S2 Fig) - this holds every year from 1975 until today, and represents a coverage about twice as high as in the gene field.

### S2 Appendix. The chemicals network

The research on chemicals and drugs is located in between the gene and the disease field with a total of 740,000 papers (S1 Table) and close to 30,000 different keywords. Here, we were most surprised by some recent developments in keyword use.

Remarkably, it is showing a distinctly dropping rate of keyword discoveries: At the end of the 1990s, the number of chemicals introduced each year stagnates at a rate of about 750 new keywords per year (S1 Fig).From 2007 on, it starts dropping (S1 Fig), regardless of the publication rate rising up until 2013 (Fig 4): More papers seem to be containing fewer new keywords.

Additionally, the reuse of known terms is becoming less frequent, resulting in the vocabulary being hardly covered every year. During the same period, the percentage of the known keywords appearing in current publications has dropped from 30 to only 20 percent, making chemicals the category with the lowest coverage (S2 Fig).

Whether this is a result of trend change over the course of he last twenty years is left an open question here.

### S3 Appendix. Keyword rate in the year 1975

A point that we felt required further investigation is the noticeable jump or drop most of the presented images show in year 1975. At once, the number of disease occurrences almost doubles (Fig 4, and density (Fig 6 as well as clustering coefficients (Fig 7) go up.

We suspect an explanation inherent to the PubMed database: In 1975, the MEDLINE data - PubMed’s main source of paper indices - was migrated from MEDLARS, the first system to enable automated queries, to its successor MEDLARS II [46]. The new machine outperformed MEDLARS in capacity, retrieval speed, and cost efficiency, it also now contained a hierarchical keyword vocabulary. It seems likely the new potential was utilized as MEDLARS II was launched, which resulted in a jump in registered publications in that year. Especially the new keyword organisation might have been impactful: the mean number of keywords makes a leap in 1975 - with the greatest effect in the disease graph, which subsequently has an additional term per paper (S3 Fig). Further evidence is provided by the fact that the rate only slightly drops afterwards. Possibly, a number of additional terms was introduced for the new hierarchy, leading to a one-time peak for new keywords and a permanent rise in the number of annotations per paper.

We are aware that this theory leaves some open questions. It is unclear why the new system would show up this locally. Our explanation implies that in 1975, numerous papers of this particular year were added - after which the rate went back to normal. However, we could not make out another event in either biochemical science or the database’s history likely to have the observed impact.

### S4 Appendix. Example abstract and discussion of the extracted links

In the following, we discuss the quality of links formed by a sampled abstract according to the link extraction method as used for our keyword graphs. The abstract by Limmroth, May, and Diener 1999 (*PMID 10023111*) [47] is reproduced in the following:

> Vasoconstrictive agents have been widely used in the treatment of migraine. LAS had no effect on BFV. Ergotamine increased BFV in the middle cerebral artery only. No correlation was found between changes in BFV and the relief of headache. This is the first trial to compare the intravenous formulation of LAS in the treatment of migraine with another antimigraine medication and suggests that it is an effective and safe drug for the parenteral treatment of acute migraine attacks. These types of drugs have various side effects and are not suitable for many patients. Due to nausea or vomiting, nonoral treatment is often required, but only a few nonvasoconstrictive drugs exist in a parenteral form and are suitable for the treatment of acute migraine in the emergency setting. In a randomized, double-blind, crossover trial we evaluated the efficacy of 1,000 mg lysine-acetylsalicylic acid i.v. (LAS) compared to 0.5 mg ergotamine s.c. in 56 patients (112 attacks) with acute migraine. To gain further insight into the possible role of vasoconstriction, blood flow velocities (BFV) were measured in intra- and extracranial arteries using duplex sonography and transcranial Doppler sonography. Both agents were equally potent in relieving headache. Intravenous LAS resulted in a significantly faster relief and had fewer side effects.

14 keywords were indexed in the example abstract, six of which are included in the neurological disorder subset: *lysine-acetylsalicylic acid, acetylsalicylic acid, ergotamine, migraine, headache* and *nausea or vomiting*. The constructed graph therefore contains links between each of the keyword nodes, which is the desired effect here for the most part.

To begin with, two semantic relations are established: *lysine-acetylsalicylic acid* is connected to the broader *acetylsalicylic acid, migraine* to *headache*. Next, the drugs *lysine-acetylsalicylic acid, acetylsalicylic acid* and *ergotamine* are linked both to the disorders they are healing, *migraine* or *headache*, and to the mentioned side effect *nausea and vomiting*. Additionally, the drugs *(lysine-)acetylsalicylic acid* and *ergotamine* are connected to each other, which is legitimated by the similar application.

A less wanted link is formed from *migraine* and *headache* to *nausea* and *vomiting*; the abstract in fact does not relate the symptoms but describes one as a side effect of a cure for the other.

**S1 Table.** Overview of the dataset. Papers belong to a category if they contain at least one associated keyword. Thus, the same paper may occur in multiple categories.

| category | before 1947 | 1947-2017 | neuro. disease 1947-2017 |
| --- | --- | --- | --- |
| chemical | 20,582 | 11,110,408 | 738,726 |
| disease | 40,643 | 15,333,771 | 2,057,272 |
| gene | 2,629 | 4,832,824 | 402,239 |
| total | 58,144 | 20,343,787 | 2,120,227 |

